# Deep, soft, and dark sounds induce autonomous sensory meridian response

**DOI:** 10.1101/2019.12.28.889907

**Authors:** Takuya Koumura, Masashi Nakatani, Hsin-I Liao, Hirohito M. Kondo

**Affiliations:** Human Information Science Laboratory, NTT Communication Science Laboratories, NTT Corporation, Atsugi, Kanagawa 243-0198, Japan; Faculty of Environment and Information Studies, Keio University, Fujisawa, Kanagawa 252-0882 Japan; School of Psychology, Chukyo University, Nagoya, Aichi 466-8666, Japan

**Keywords:** ASMR, hearing, somatosensory, interaural difference, proximal space

## Abstract

There has been a growing interest in the autonomous sensory meridian response (ASMR). The ASMR is characterized by a tingling sensation around the scalp and neck and often induces a feeling of relaxation and a reduction of a negative mood. However, it is still unknown what factors affect the ASMR. The present study focused on stimulus characteristics and individuals’ mood states and personality traits. Participants filled out self-reported questionnaires (the Profile of Mood States, Beck Depression Inventory, and Big Five Inventory) and reported ASMR estimates throughout a 17-min experiment while listening to binaural tapping and brushing sounds. Cross-correlation results showed that the ASMR estimates were strongly associated with the acoustic features of auditory stimuli, such as their amplitude, spectral centroid, and spectral bandwidth. This indicates that low-pitched sounds with dark timbre trigger the ASMR. The maximum ASMR was observed around 2 s after the acoustic features changed, suggesting that the sluggishness of multisensory integration may lead to the ASMR experience. In addition, individual differences in the ASMR experience were closely linked to participants’ mood states, such as anxiety, but not to their personality traits. Our results provide important clues to understand the mechanisms of auditory-somatosensory interactions.

**Significant Statements:** The autonomous sensory meridian response (ASMR) is characterized by a tingling, electrostatic-like sensation across the scalp and back of the neck. This phenomenon can be triggered by a variety of audiovisual stimuli, and many people seek out the ASMR via the internet to receive a feeling of relaxation and reduce a negative mood. We show that the ASMR is induced about 2 s after acoustic features, such as the amplitude, spectral centroid, and spectral bandwidth are changed. This suggests that low-pitched sounds with dark timbre lead to the ASMR experience. The stimulus-driven ASMR effect is found regardless of the personality traits or mood states of participants. Our findings provide a critical clue to understand the mechanisms of auditory–somatosensory interactions.

## Introduction

Many regular-to-heavy users of the internet are familiar with the autonomous sensory meridian response (ASMR), a phenomenon often characterized by a tingling, electrostatic-like sensation across the scalp and back of the neck (Barratt and Davis, 2015). As the ASMR can be triggered by a variety of audiovisual stimuli, content creators have produced hundreds of ASMR-inducing videos for video-sharing platforms such as YouTube. Many individuals experience ASMR to receive a feeling of relaxation, reduce a negative mood, and aid sleep (Barratt and Davis, 2015).

In contrast to the general population’s interest in ASMR, few scientific studies sought to clarify the types of stimuli that most effectively trigger the ASMR and how the ASMR is related to an individual’s mental states and personality traits. Recent studies have used web-based surveys of the Big Five Inventory (BFI) and demonstrated that individuals with higher Openness and lower Conscientiousness scores reported more frequent ASMR experiences (McErlean and Banissy, 2017; Fredborg et al., 2018). However, such studies include an inherent sample selection bias in that participants recruited through the internet may have the tendency to disclose their ASMR experiences. In addition, how the stimulus characteristics were controlled is unclear, which may inherently affect the ASMR.

From the perspective of cross-modal interactions, the ASMR is an intriguing phenomenon because sounds induce a shiver or thrill that corresponds to a tickling sensation on the skin (Grewe et al., 2009). There is no doubt that the integration of vision, touch, and proprioception plays a critical role in shaping the body schema (Maravita et al., 2003; de Vignemont et al., 2005; Haggard et al., 2007). However, few studies have looked at the contribution of audition to the somatosensory system. Effects of auditory distractors on tactile discrimination are larger when the distractors are presented close to the head (20 cm) than when they are presented far from the head (70 cm) (Kitagawa et al., 2005). Self-produced action sounds bias the body schema and its spatial boundaries (Tajadura-Jimenez et al., 2012). In addition, auditory feedback can modify the tactile sensation of a listener experiences when rubbing his/her hands together (Jousmäki and Hari, 1998; Guest et al., 2002). When high-frequency components of the auditory feedback increase, the perceived roughness/moisture of the palmar skin decreases and its smoothness/dryness increases. Actually, listeners reported a tickling sensation when listening to sounds while the ear of a dummy head was stroked with a brush (Kitagawa and Igarashi, 2005). However, it is unclear what factors transform auditory inputs into the tickling sensation.

The present study examined the degree to which the ASMR is affected by the acoustic features of auditory stimuli, as well as by mood states and personality traits of individuals. Participants, all of whom had never experienced the ASMR, were instructed to listen to binaural tapping and brushing sounds from headphones. We hypothesized that there should be no confounding factor between acoustic features and an individual’s states/traits because these parameters are essentially independent of each other. The ASMR may be similar to aesthetic chills in terms of a sound-induced emotional experience. After the experiment, we interviewed listeners to investigate the pleasantness of the ASMR experience.

## Materials and Methods

### Listeners

Thirty college students (13 males and 17 females; mean age 20.6 years, range 18–23 years) participated in this study. Listeners were right-handed with normal hearing. None had ever viewed any video created to induce the ASMR experience. All listeners gave written informed consent after the procedures had been fully explained to them. The procedures reported in this study were approved by the Ethics Committee of School of Psychology at Chukyo University (protocol no.RS17-020).

### Behavioral tasks

Auditory stimuli were made from a file with Creative Commons licenses that was produced to induce the ASMR and uploaded to the video-sharing site YouTube (https://www.youtube.com/watch?v=7MZtaAgqoTY). The total duration of each auditory stimulus was 1,020 s, and visual information was removed from it. Listeners were instructed to dichotically listen to the stimulus through headphones (Sennheiser HD 599) and continuously indicate their sensation using a 3-point Likert scale: “no ASMR”, “weak ASMR”, or “strong ASMR”. In this study, ASMR was defined as a tingling sensation with “zoku-zoku”, which is a Japanese onomatopoeia word for shivering with a feeling of being excited and frightened (Osaka et al., 2003; Osaka et al., 2004). Their responses were collected via three keys on a computer keyboard with the sampling rate of 1 kHz. A key press indicating a response was held until a subsequent key press. The stimulus presentation and data collection were controlled by a PC with Presentation software (Neurobehavioral Systems, Berkeley, CA, USA). We interviewed listeners separately after the experiment. In the interviews, they reported where their tingling sensation originated and whether the sensation moved to other areas of their body. In addition, we checked the degree of their ticking sensation and pleasant feeling using a visual analogue scale.

### Questionnaires of mood states and personality traits

Before the experiment, listeners filled out questionnaires in a quiet room. We obtained self-reported measures of listener’s mood states and personality traits using the POMS, BDI-II, and the BFI. The POMS measures transient, distinct mood states for the past one week and consists of six subscales: Tension-Anxiety (9 items), Depression (15 items), Anger-Hostility (12 items), Vigor (8 items), Fatigue (7 items), and Confusion (7 items) (McNair et al., 1971). Listeners rated their mood by 65 adjectives (e.g., “Furious”, “Hopeless” and “Carefree”) on a 5-point Likert scale from “not at all” to “extremely”. The BDI is widely used as an indicator of the severity of depression in adolescents and adults (Beck et al., 1996). Listeners responded to 21 items based on their mental states over the past two weeks. The BFI is used to measure the individual’s basic personality traits: Extraversion (8 items), Agreeableness (9 items), Conscientiousness (9 items), Neurotisism (8 items), and Openness (10 items) (John and Srivastava, 2001; Kondo et al., 2017). Listeners rated these 44 items on a 5-point Likert scale ranging from “strongly disagree” to “strongly agree”. One listener was excluded from the subsequent analysis on mood states and personality traits because his/her data on the questionnaires were missing.

### Data analyses

#### ASMR data analysis

The time-series data of the ASMR estimates were converted to numerical values of 0, 1, and 2 (Fig. 1). Based on the interviews with participants, the ASMR-induced tingling sensation was visualized on areas of the body (Fig. 2). We performed a correlation analysis to check the relationships between averaged ASMR estimates, mood states, and personality traits, and calculated a significance level. Statistical analyses were carried out with Python, R (version 3.1.2), and IBM SPSS Statistics (version 23).

**Figure 1.**
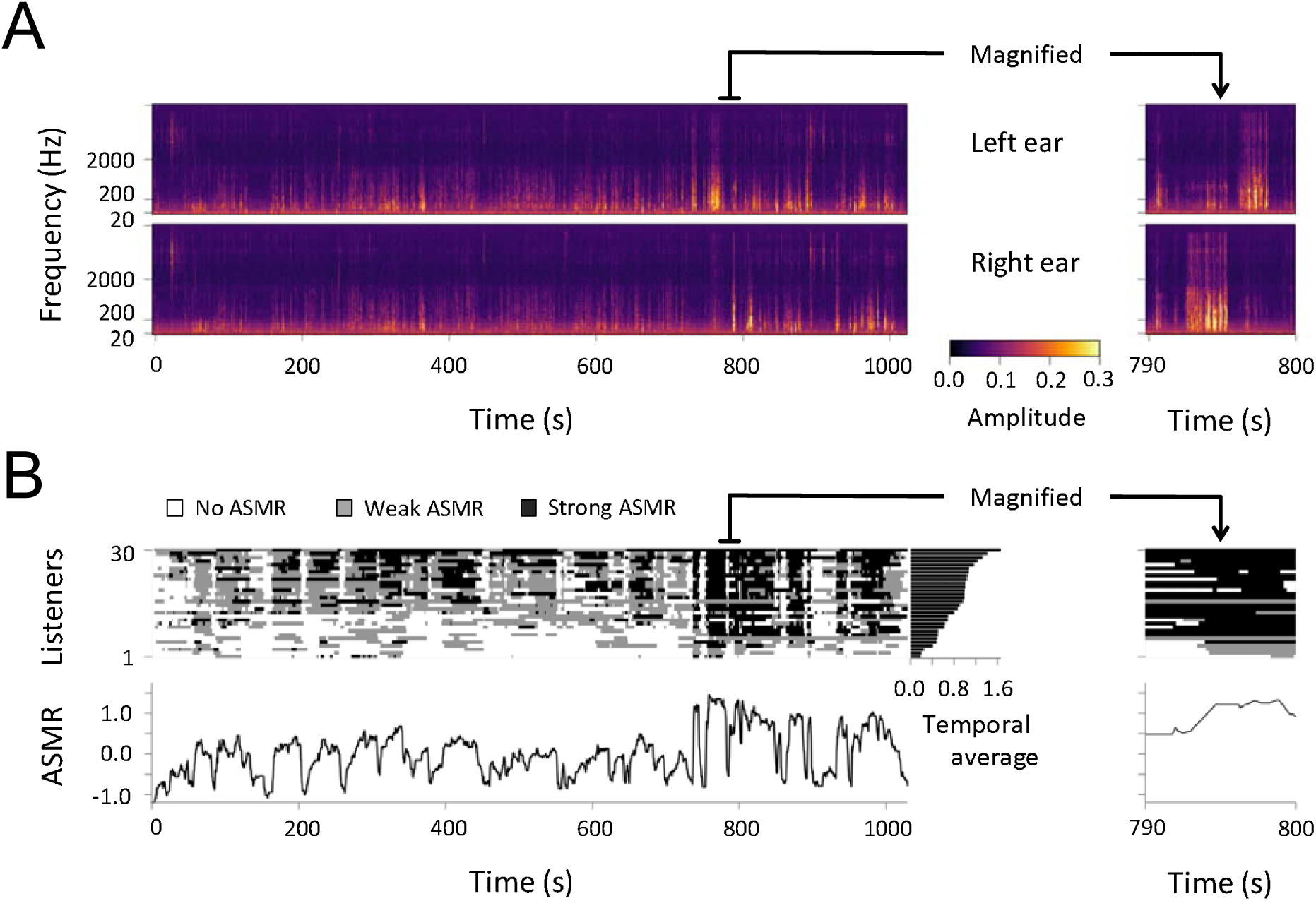
Auditory stimuli and ASMR estimates. (A) Cochlea spectrograms of the stimuli recorded by the artificial ear. (B) Results of the ASMR estimates. Top and bottom panels represent time-series data for each listener and averaged *z*-scores across the listeners, respectively. The right panel represents temporal average in each listener.

**Figure 2.**
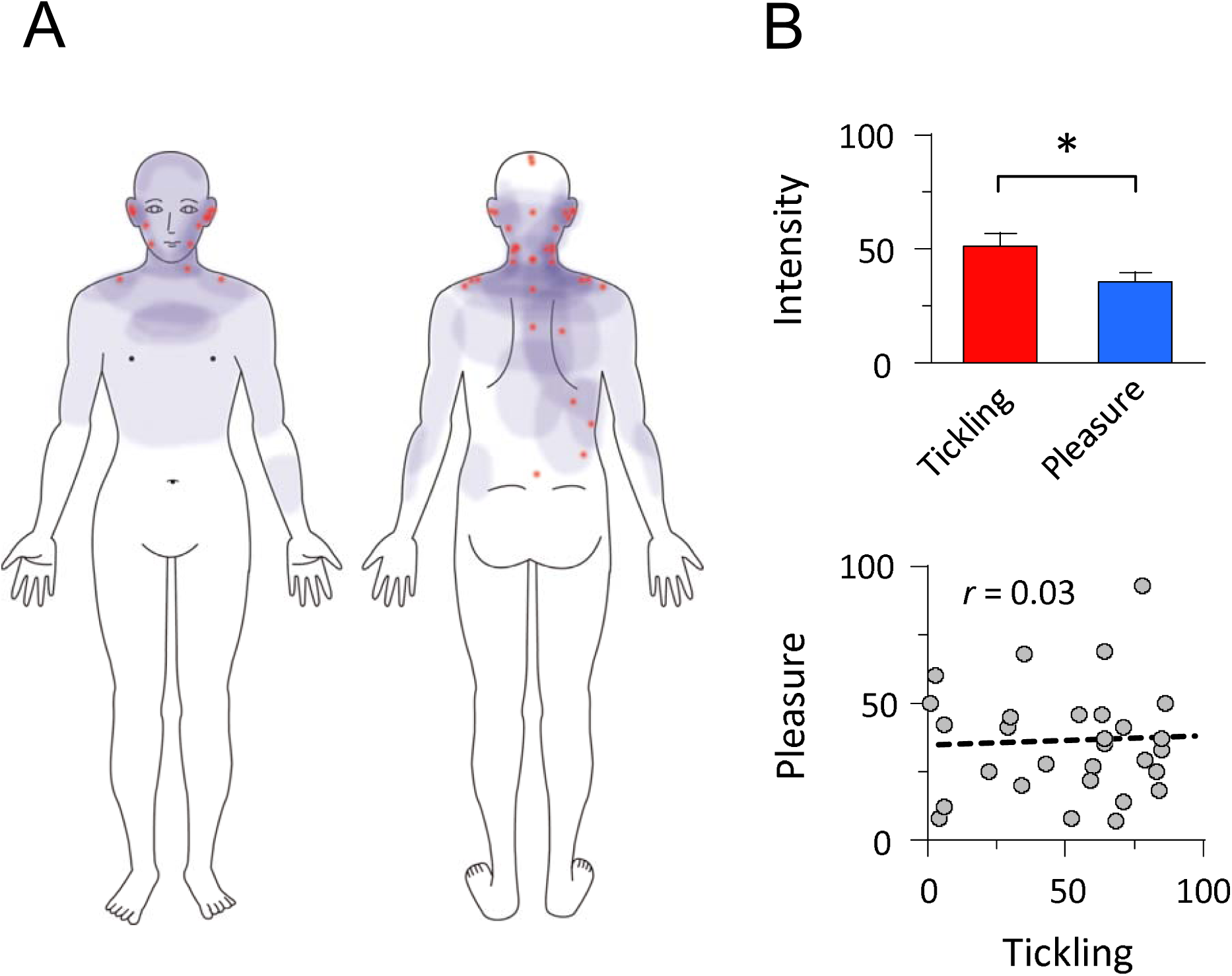
ASMR-induced tingling sensation and pleasant feeling. (a) An illustration of tickling sensation. Red spots indicate an origin of the sensation, whereas areas colored in pale blue represent the sensation spreading to other parts of the body. (b) The relationship between tickling sensation and pleasant feeling. Bars represent the means of subjective rating, whereas circles indicate individual data. Error bars represent the standard errors of means. The dashed line represents a linear regression. \**p* < 0.05.

#### Acoustic features of auditory inputs

The cochlear spectrogram was calculated from the auditory stimuli presented to each ear. The sounds were recorded with an artificial ear (TYPE2015, ACO, Tokyo) with the sampling rate of 44.1 kHz. The recorded waveforms were decomposed into frequency components by a gammatone filter bank (Patterson et al., 1992). The center frequencies of the filters were spaced one equivalent rectangular bandwidth (ERB) apart (Glasberg and Moore, 1990), and the lowest center frequency was set to 20 Hz, resulting in 42 filters with the highest center frequency of 20.562 kHz. To simulate non-linear compression in the auditory periphery, amplitude envelopes of the frequency-decomposed signals were calculated with Hilbert transformation, powered by 0.3 (McDermott and Simoncelli, 2011), and down-sampled to 2 kHz.

The auditory features obtained for both the left and right channels were the amplitude, spectral centroid, spectral bandwidth, and instantaneous roughness. The amplitude is the mean of the cochlear spectrogram over frequencies:

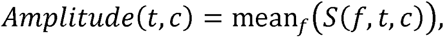

where *f, t*, and *c* denote a frequency bin in the ERB scale, time, and a channel indicating left or right, respectively. *S*(*f, t, c*) represents a cochlear spectrogram. The spectral centroid indicates the gravity center in the frequency domain:

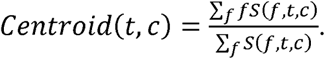

The spectral bandwidth is the width of the mass distribution in the frequency domain:

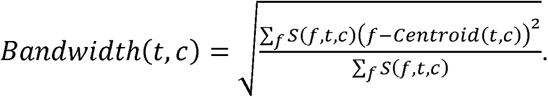

Roughness is often calculated from a sound segment with a certain duration. To evaluate temporal variation of the stimulus, we calculated roughness at every time point, that is, an instantaneous roughness. For the instantaneous roughness, the cochlear spectrogram at each center frequency was filtered with a bandpass filter with a passband frequency between 30 and 150 Hz (Arnal et al., 2015). Instantaneous roughness was defined as

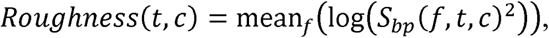

where *S_bp_* denotes the bandpass-filtered spectrogram.

The four features described above were down-sampled to 10 Hz. We computed the average and the absolute difference of the features in the left and the right ears.

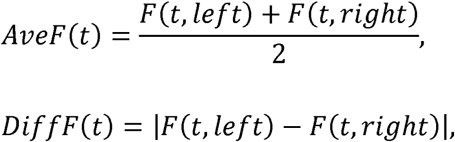

where *F* indicates amplitude, centroid, bandwidth, or roughness. This resulted in totally 8 features = 4 (amplitude, spectral centroid, spectral bandwidth, and instantaneous roughness) × 2 (average and absolute difference). We refer to them as Ave- and Diff-amplitude, Ave- and Diff-centroid, Ave- and Diff-bandwidth, and Ave- and Diff-roughness.

#### Correlations between ASMR and acoustic features

We calculated cross-correlations between the time course of the ASMR estimates and acoustic features. The ASMR estimates were *z*-scored for each listener and averaged over listeners. The correlations were assessed by Spearman’s rank-order correlations. Statistical significance of the correlation coefficient at the peak of the cross-correlogram was tested by comparing the peak correlation coefficient with 16,000 cross-correlogram calculated from randomly shifted time courses. We also calculated a cross-correlogram for each listener. The cross-correlations did not reach statistical significance for one listener on the amplitude; for two listeners on the spectral centroid; and for five listeners on the spectral bandwidth. Consequently, we excluded the data of six listeners from a repeated-measures ANOVA on the time lags of cross-correlations when comparing between the acoustic features (*N* = 24).

#### Relationship between ASMR and individual’s states and traits

For the factor analysis, we checked the Kaiser–Meyer–Olkin (KMO) statistic using zero-order correlations and partial correlations to test whether the variables in our dataset were adequate to correlate. The KMO statistic (0.67) indicated that some common factor underlay our dataset, because its estimate was higher than 0.50 for proceeding with a satisfactory factor analysis. Bartlett’s test of sphericity showed that correlations between the variables were greater than those expected by chance: *χ*^2^ = 255.04, *p* < 0.001. The factors were extracted by least squares estimation and then subjected to an oblique promax rotation.

## Results

### ASMR experience

Time-series data of auditory stimuli and ASMR estimates are shown in Figure 1. Tapping and brushing sounds were used as auditory stimuli to induce the ASMR. The results show not only interindividual variations of the ASMR estimates but also similar temporal changes across participants. After the experiment, ninety percent of the participants reported an ASMR-induced tingling sensation on different parts of their body (Fig. 2a). Of those parts, it mainly originated on the ears and their vicinities (59%), the neck (44%), the shoulders (44%), and the spine and back (30%). Our results are consistent with previous findings on the ASMR experience (Barratt and Davis, 2015). A previous study on music-induced chills found that most participants experienced goosebumps on their arms, whereas less than half felt anything in their spine (Craig, 2005). In this study, however, small numbers of participants reported the tingling sensation on their arms (15%) and legs (0%). Using a visual analogue scale (range 0 to 100), we conducted an experiment to check the subjective intensity of participants’ tingling and pleasure (Fig. 2b). The intensity (mean ± SE) was greater for tickling (51.2 ± 5.3) than for pleasure (35.7 ± 3.7): *t* = 2.43, Cohen’s *d* = 0.63, *p* = 0.022. There was no significant correlation between the intensity of the two measures: *r* = 0.03, *p* > 0.87. In addition, the ASMR estimates were positively correlated with the intensity of the tingling sensation (*r* = 0.74, *p* < 0.001), but not with the intensity of pleasantness (*r* = 0.00, *p* > 0.98). Thus, the results suggest that the ASMR is not necessary linked with pleasant experiences, at least not for the stimuli used in the present study.

### Correlations between ASMR and acoustic features

We performed a cross-correlation analysis to examine whether and how each acoustic feature contributed to the ASMR as a function of time (Fig. 3). The acoustic features we examined were the amplitude, spectral centroid, spectral bandwidth, and instantaneous roughness. We computed an average (Ave) of the features presented to the left and right ears and the difference (Diff) in the features between the two ears. For the Ave-features, the ASMR estimates were positively correlated with the Ave-amplitude (*r*_*s*_ = 0.37, *p* < 0.001), indicating that auditory stimuli with larger sound pressure levels induce greater ASMR estimates (Fig. 3). The ASMR estimates were negatively correlated with the Ave-centroid (*r*_*s*_ = −0.48, *p* < 0.001) and Ave-bandwidth (*r*_*s*_ = −0.34, *p* < 0.001). This suggests that low-pitched sounds with dark timbre (a centroid frequency of less than 1.5 kHz) lead to an ASMR experience. In contrast, the correlation between the ASMR estimates and Ave-roughness did not reach statistical significance (*r*_s_ = 0.24, *p* = 0.088).

**Figure 3.**
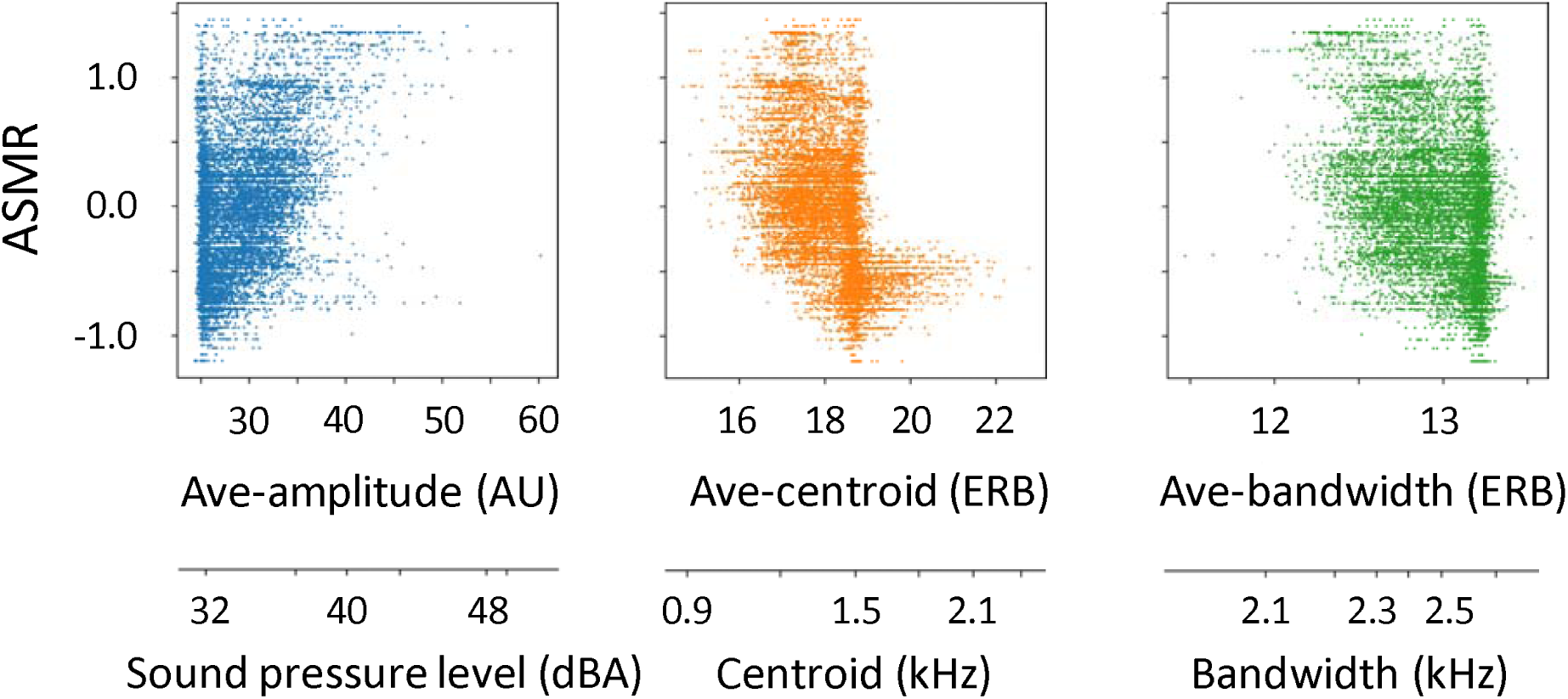
Results of correlation analysis on the ASMR estimates. Plots of ASMR estimates (*z*-scores) and each acoustic feature. ASMR estimates are shifted by the peak lags of cross-correlations. When the Ave-amplitude was larger, the ASMR estimate was greater. In contrast, when Ave-centroid and Ave-bandwidth were smaller, it was greater. Average values of corresponding physical dimensions are shown below the panels, although they do not have a one-to-one correspondence to acoustic feature values. AU: arbitrary unit. ERB: equivalent rectangular bandwidth scale.

We also found significant cross-correlations between the ASMR estimates and all the Diff-features except for instantaneous roughness: *r*_*s*_ = 0.28, *p* < 0.001 for the Diff-amplitude; *r*_*s*_ = 0.25, *p* < 0.001 for the Diff-centroid; *r*_*s*_ = 0.24, *p* < 0.001 for the Diff-bandwidth; cf., *r*_*s*_ = 0.20, *p* = 0.053 for the Diff-roughness (Fig. 4). This means that as the larger the interaural differences in the four acoustic features were, the larger the ASMR estimates became. Intriguingly, the peak of cross-correlation coefficients had a time lag of around 2 s. These results indicate that changes in the acoustic features of auditory stimuli trigger the ASMR.

**Figure 4.**
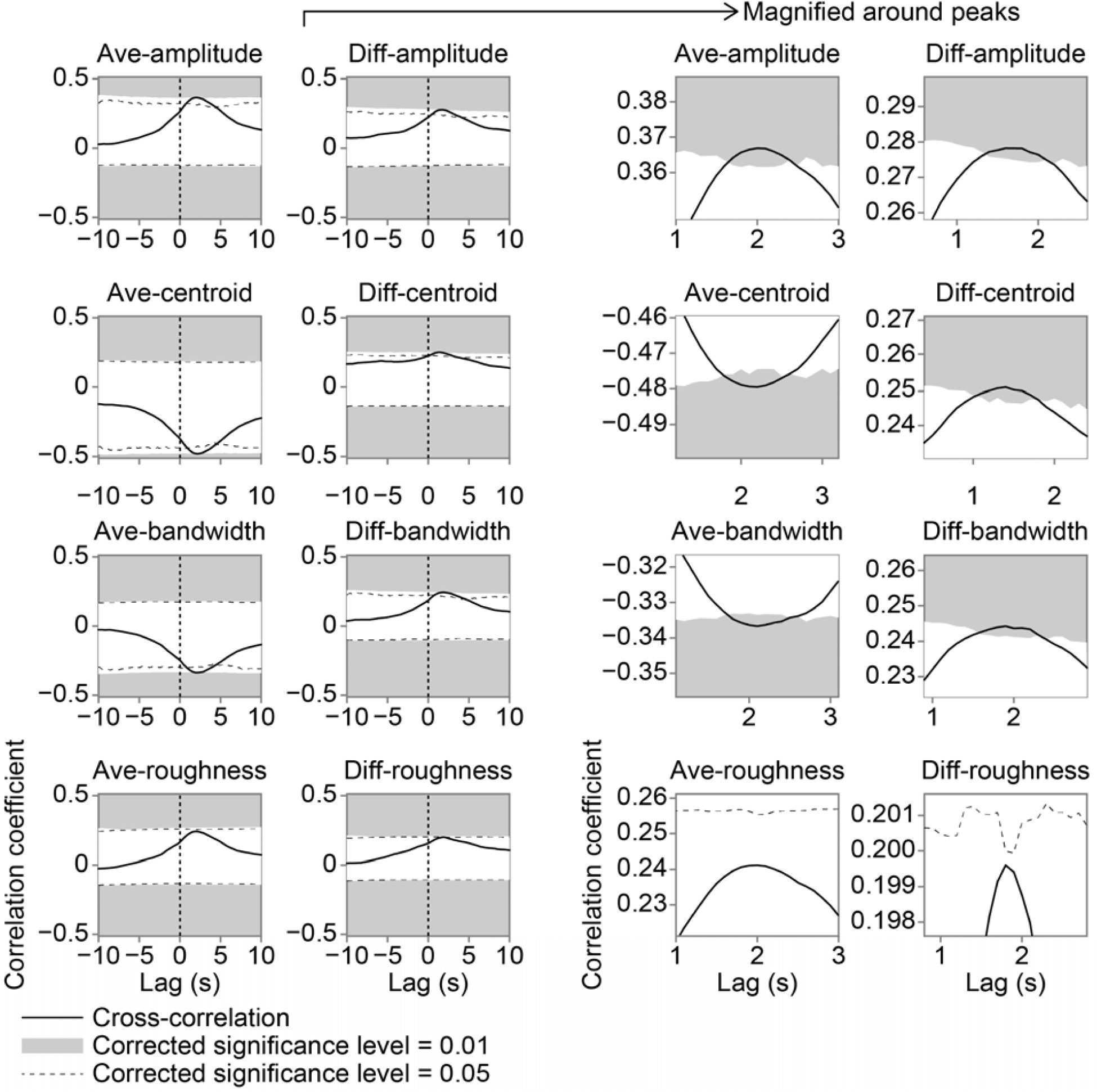
Cross-correlations between the ASMR estimates and acoustic features. Areas shaded in grey and dashed lines represent statistical significance after Bonferroni correction over the eight features. The right panels show magnified plots around peaks of cross-correlations. The positive time lag indicates that the changes in the acoustic features are followed by the ASMR.

We further examined the time lag of cross-correlations using the individual differences approach. The Ave- and Diff-roughness features were excluded from the subsequent analyses because their cross-correlations did not reach statistical significance. We performed a 2 (Ave and Diff) × 3 (amplitude, centroid, and bandwidth) analysis of variance (ANOVA) on the time lags to the peak of cross-correlations (Fig. 5). The time lag of Diff-features (1.78 ± 0.17 s) was shorter than that of Ave-features (2.22 ± 0.19 s): 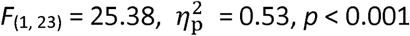. The main effect ofthe acoustic features did not reach statistical significance—1.93 ± 0.15 s for amplitude; 1.81 ± 0.14 s for centroid; 2.17 ± 0.26 s for bandwidth; 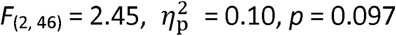—and neither did the interaction: 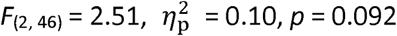. Therefore, our results suggest that the ASMR is caused by interaural differences in the acoustic features and consequently affected by the average of those features.

**Figure 5.**
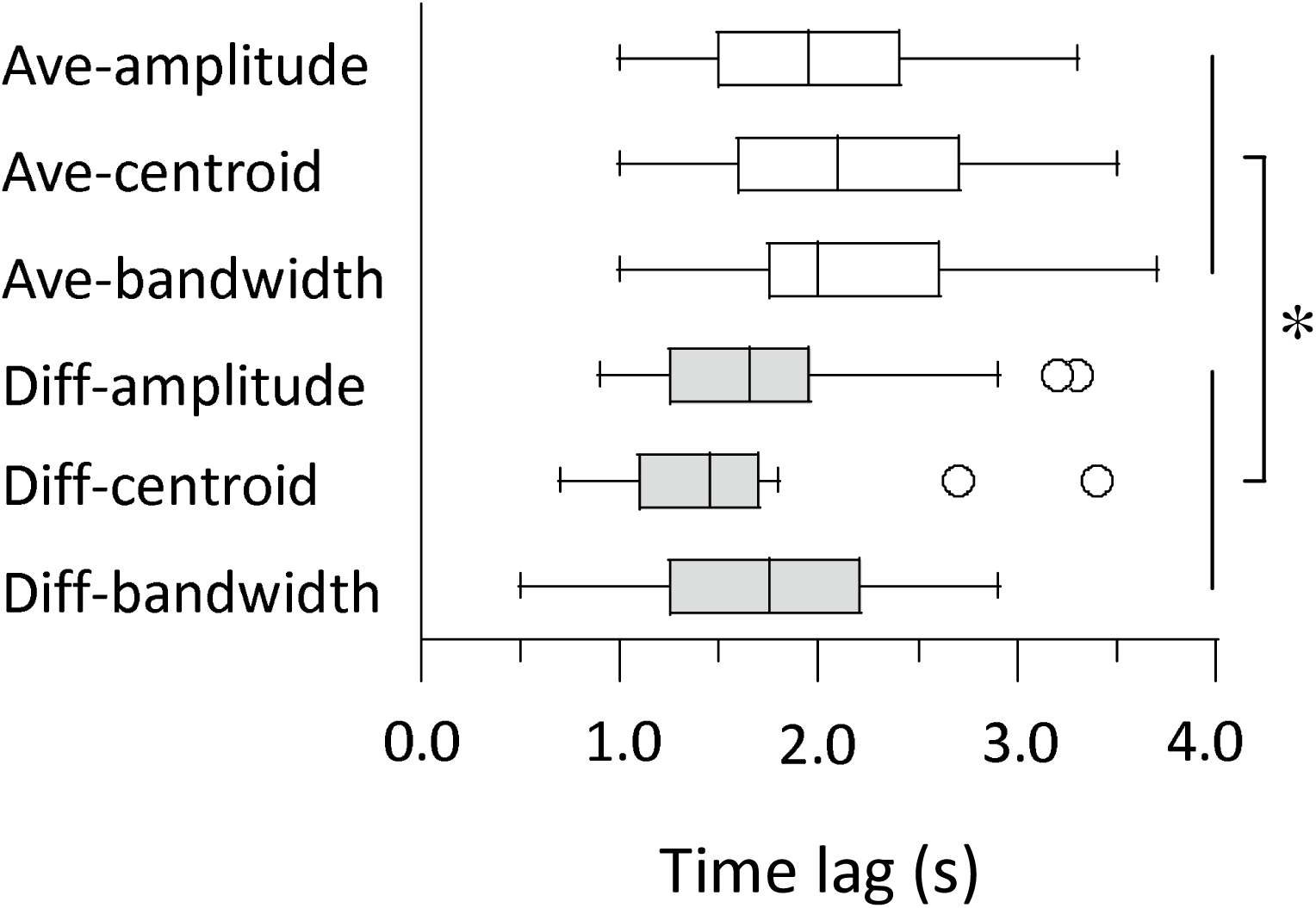
Time lags of peak cross-correlations for the acoustic features. Box-and-whisker plots indicate the median, 25th and 75th percentiles, and outliers for the distribution across listeners. \**p* < 0.05.

### Effects of states and traits on ASMR

So far, we have demonstrated effects of acoustic features on the ASMR. Next, we investigated whether the ASMR is associated with the mood states and personality traits of the listeners, which were assessed using the Profile of Mood States (POMS), Beck Depression Inventory (BDI-II), and the BFI (Table 1). Cronbach’s alpha of the measures reached a satisfactory level (range of 0.65 to 0.93), suggesting that these measures have a high level of internal consistency. The ASMR estimates were significantly correlated with the POMS Anxiety score: *r* = 0.52, *p* = 0.004 (Table 2). There were mild positive correlations (0.42 to 0.46) between the ASMR estimates and POMS Fatigue, Depression, and Confusion scores. In contrast, the correlations with the BFI scores were generally low (less than 0.32). The overall results indicate that individual’s mood states, relative to his/her personality traits, are involved in the ASMR experience. We used a multiple regression analysis to specify the degree to which the twelve states/traits scores contribute to the ASMR estimates. The results of a stepwise analysis showed that the POMS Anxiety score accounted for 24.5% of the variance of the ASMR estimates: adjusted *R*^2^ = 0.208, *F*_(1, 20)_ = 6.50, *p* = 0.02. However, the contribution of the other measures was not significant. Thus, it is likely that individuals with anxiety or depressive states are sensitive to the ASMR.

**Table 1.** Descriptive statistics for ASMR estimates and individual’s states/traits.

| Measure | Mean | SD | Min | Max | Skewness | Kurtosis | Reliability |
| --- | --- | --- | --- | --- | --- | --- | --- |
| ASMR | 0.84 | 0.36 | 0.19 | 1.66 | -0.01 | -0.33 | N/A |
| POMS |  |  |  |  |  |  |  |
| Tension-Anxiety | 13.5 | 6.9 | 1 | 31 | 0.43 | 0.45 | 0.82 |
| Depression | 16.4 | 12.3 | 0 | 46 | 0.65 | 0.02 | 0.93 |
| Anger-Hostility | 11.3 | 9.6 | 0 | 34 | 0.75 | -0.25 | 0.93 |
| Vigor | 10.6 | 6.7 | 0 | 30 | 0.88 | 1.20 | 0.91 |
| Fatigue | 11.9 | 6.0 | 2 | 27 | 0.45 | 0.26 | 0.86 |
| Confusion | 11.4 | 6.1 | 0 | 25 | 0.01 | -0.03 | 0.84 |
| BDI-II | 12.7 | 8.1 | 2 | 31 | 0.67 | -0.12 | 0.81 |
| BFI |  |  |  |  |  |  |  |
| Extraversion | 21.2 | 6.0 | 10 | 34 | 0.24 | -0.63 | 0.80 |
| Agreeableness | 28.0 | 6.2 | 10 | 38 | -0.78 | 1.23 | 0.65 |
| Conscientiousness | 26.2 | 5.9 | 13 | 36 | -0.32 | -0.42 | 0.76 |
| Neuroticism | 26.7 | 6.0 | 9 | 35 | -1.29 | 1.82 | 0.80 |
| Openness | 29.5 | 7.0 | 16 | 41 | -0.13 | -0.95 | 0.76 |

Reliability was calculated using Cronbach's alpha. ASMR, autonomous sensory meridian response; BDI, Beck Depression Inventory; BFI, Big Five Inventory; POMS, Profile of Mood States.

**Table 2.** Correlations between ASMR estimates and individual’s states/traits.

| Measure | Correlation |  |
| --- | --- | --- |
|  | *r* | *p* |
| POMS |  |  |
| Tension-Anxiety | **.521** | **.004** |
| Depression | .420 | .023 |
| Anger-Hostility | .267 | .162 |
| Vigor | -.141 | .465 |
| Fatigue | .455 | .013 |
| Confusion | .416 | .025 |
| BDI-II | .281 | .140 |
| BFI |  |  |
| Extraversion | -.101 | .595 |
| Agreeableness | .093 | .623 |
| Conscientiousness | .135 | .476 |
| Neuroticism | .318 | .087 |
| Openness | -.266 | .155 |

A value in bold font is significant after a false discovery rate (FDR) correction ( $N = 29$ ).

We performed a factor analysis to further examine the relationships between the ASMR estimates and states/traits scores. The BFI Agreeableness score was excluded from the subsequent analyses because it was closely related to the BFI Conscientiousness score (*r* = 0.51, *p* = 0.005) and its reliability was relatively low (0.65). Two factors, whose eigenvalues were larger than 1, were extracted from the 12 subscales (Fig. 6). The first factor, with an eigenvalue of 5.78 before the promax rotation, was heavily loaded on the ASMR estimates, all the POMS subscores except the Vigor score, the BDI score, and the BFI Neuroticism score (factor loadings, 0.48 to 0.93). The second factor, with an eigenvalue of 1.68, was loaded on the BFI Extraversion and Openness scores and the POMS Vigor score (0.49 to 0.97). The first and second factors were termed the “anxiety” and “activity”, respectively. The contribution of the “anxiety” factor (0.51) to the ASMR estimates was much greater than that of the “activity” factor (0.05).

**Figure 6.**
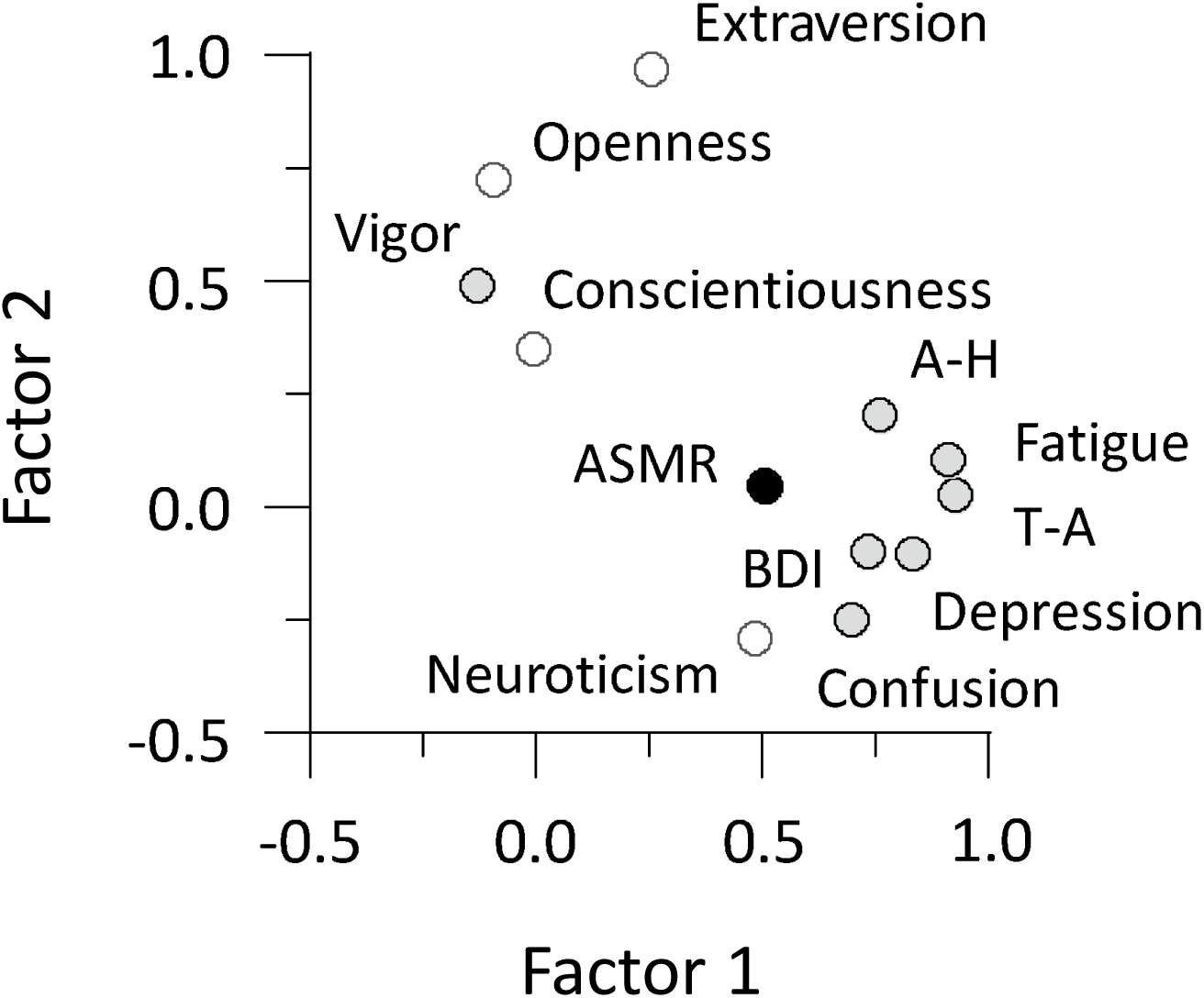
Results of the factor analysis. The filled and open circles indicate the subscales of questionnaires on individual’s mood states and personality traits, respectively. Factors 1 and 2 can be considered the “anxiety” and ‘‘activity’’ factors. A-H: Anger-Hostility. T-A: Tension-Anxiety.

## Discussion

The present results demonstrate that the intensity of ASMR estimates varied across listeners, although its sensation mainly originated on their ears, neck, and shoulders. The ASMR was closely linked to the tickling sensation but not to the pleasant-unpleasant dimension. The ASMR was affected by the acoustic features of auditory stimuli. Specifically, the cross-correlation results showed that the ASMR was induced by interaural differences in the acoustic features, such as the amplitude, spectral centroid, and spectral bandwidth. The maximum ASMR was observed around 2 s after the acoustic features changed. In addition, it was associated with listener’s mood states, including anxiety. In what follows, we mainly discuss effects of stimulus characteristics on the ASMR.

Many internet users expect the ASMR to provide a feeling of relaxation (Barratt and Davis, 2015). Our preliminary study using a test-retest paradigm indicated that the ASMR estimates have high reliability within ASMR-inexperienced participants: *r* = 0.87, *p* = 0.001, *N* = 10. It seems that the ASMR differs from music-induced chills in terms of physiological, emotional, and phenomenological aspects. Piloerection during chills is innervated by the sympathetic nervous system (Lindsley and Sassaman, 1938; Craig, 2005). There is no consensus that the parasympathetic nervous system is involved in the ASMR (Poerio et al., 2018). A profound emotional experience, such as weeping, sometimes arises in response to music-induced chills (Sloboda, 1991; Panksepp, 1995; Mori and Iwanaga, 2017), but not in response to the ASMR. In contrast to aesthetic chills, the ASMR is frequently triggered by a variety of stimuli, such as whispering, close personal attention, and crisp sounds (Barratt and Davis, 2015). The present study investigated common factors (i.e., acoustic features) affecting various ASMR stimuli.

We have demonstrated that the Diff-features of auditory stimuli play an important role in triggering the ASMR. Specifically, a larger Diff-amplitude parameter resulted in larger ASMR estimates. The Diff-amplitude used in the present study corresponds to an interaural level difference (ILD). Thus, the results can be interpreted as indicating that the ASMR is elicited when listeners perceive sound sources near their ears. This is consistent with the findings that the ASMR is longer during binaural listening than during diotic listening (Liao et al., 2017) and stronger for sounds moving around the head than for static sounds (Honda et al., 2019). In a previous study, listeners reported a stronger tickling sensation when the sound of an ear being tickled was presented with headphones than when the sound was presented via a loudspeaker placed 80 cm from their ears (Kitagawa and Igarashi, 2005). Therefore, it is likely that auditory–somatosensory interactions observed as the ASMR are related to sound localization ability in the space close to the head.

Most sound localization studies have focused on auditory events in the distal region (i.e., distances of more than 1 m from listeners) (Brungart and Rabinowitz, 1999). This may be because the detection of distant events in an environment is believed to be necessary for the survival of humans and mammals (Heffner and Heffner, 1992; Kitagawa and Spence, 2006). In general, the ILD and interaural time difference (ITD) are used as localization cues, respectively, when high- and low-frequency sounds reach the ears (Rayleigh, 1907; Moore, 2013). ILDs can be accurately obtained at high frequencies (more than 1.5 kHz) because diffraction due to the head hardly occurs, whereas ITDs can be easily detected at low frequencies because phase ambiguity hardly occurs. Intriguingly, a previous study demonstrated that ILDs, even at low frequencies, increase when a sound source is located less than 50 cm from the head (Brungart and Rabinowitz, 1999).Thus, we believe that listeners in our experiment used ILDs, as well as ITDs, to estimate the distance and direction of a sound source, which results in the enhancement of sensitivity to the ASMR.

Our results showed that the Ave-centroid feature was negatively correlated with the ASMR estimate. The spectral centroid has been shown to be correlated with the perceptual brightness of musical sounds with clear pitch (Almeida et al., 2017). Although our stimuli were non-musical natural sounds, a previous study revealed that both pitched and unpitched sounds induce timbre in a common perceptual space, which includes the spectral centroid as one of its major dimensions (Lakatos, 2000). Thus, it is likely that sounds with a dark impression are closely linked to the ASMR experience.

In contrast, the Ave- and Diff-roughness had a limited effect on the ASMR. It is known that roughness in sounds such as screams induces unpleasant perception (Arnal et al., 2015). The perceptual attribute can be differentiated from other communication signals such as speech. In this study, the cross-correlation between ASMR estimates and Ave-roughness was not significant but was positive. This is consistent with the behavioral results indicating that the ASMR is not necessarily a pleasant experience for some listeners. Alternatively, listeners might experience an unpleasant feeling because the auditory stimuli were not accompanied with visual information. When visual cues are not available, interactions between auditory and somatosensory information occur easily in the space behind the back (Kóbor et al., 2006; Zampini et al., 2007). It has been believed that many people expect to receive a feeling of relaxation and/or pleasure from the ASMR. Thus, a future study should investigate the relationship between the ASMR and unpleasant feelings.

It may be hypothesized that a sluggish integration of multisensory information, in particular between audition and tactile sensation, leads to the ASMR. Whether the tickling sensation occurs or not appears to depend on comparing an internal prediction with actual sensory feedback, resulting in the phenomenon where a self-produced tactile sensation differs from an externally produced one (Blakemore et al., 1998). In the present study, the maximum ASMR occurred around 2 s the Diff-features had been changed. Previous studies using auditory streaming showed that effects of head movement on perceptual formation are maximized 1.8 s after motion onsets (Kondo et al., 2012; Kondo et al., 2014). Intriguingly, the time scale of auditory–somatosensory interactions is similar to that of auditory–motor interactions. Thus, there is the possibility that the tickling sensation accompanying the ASMR is derived from the sluggishness of multisensory integration processes.

The auditory and somatosensory systems have similar temporal frequency channels (Yau et al., 2009; Occelli et al., 2011). Many neuropsychological studies have investigated somatosensory responses in the auditory cortex. Neuronal responses in the caudomedial area of the auditory cortex have been elicited by cutaneous stimulation on the neck and hands of monkeys (Schroeder et al., 2001; Fu et al., 2003). Tactile stimulation has been shown to enhance the activity of posterior auditory areas (Foxe et al., 2002; Kayser et al., 2005; Schürmann et al., 2006). In contrast, little is known about how neurons in the cerebral cortex encode information on auditory distance. A substantial study demonstrated that tactile-sensitive neurons in the ventral premotor area respond to auditory stimuli presented within 30 cm from the head (Graziano et al., 1999) and that neural responses decrease when they are presented at distances of 50 cm from the head. The traditional view on sensory systems is that different sensory information is separately analyzed in unisensory areas and subsequently projected into multisensory areas (Felleman and Van Essen, 1991). However, recent fMRI studies have revealed that auditory stimulation influences the somatosensory area (Liang et al., 2013; Pérez-Bellido et al., 2018). Thus, the present findings on the ASMR could contribute to clarifying neural mechanisms of auditory–somatosensory integration.

The results of a factor analysis showed that listeners with a higher anxiety state were more sensitive to the ASMR. An early study also reported that ASMR media improve participant’s mood, with the positive effect lasting several hours (Barratt and Davis, 2015). In contrast to previous findings (McErlean and Banissy, 2017; Fredborg et al., 2018), the ASMR estimates were not correlated with listener’s personality traits. The discrepancy in these results may be due to our using ASMR-inexperienced listeners in this study. Participants’ characteristics and learning effects probably shape the preference for ASMR media.

Previous studies have focused on the relationship between the ASMR experience and an individual’s personality traits, but have not paid attention to effects of sensory input itself on the ASMR. An essential aspect of the ASMR is that auditory stimuli produce different forms of perception at the surface of the body or inside it. The present study demonstrated that the ASMR is induced by changes in interaural differences in the acoustic features, namely the amplitude, spectral centroid, and spectral bandwidth. We pointed out that the sluggishness of multisensory integration may lead to ASMR. We also found that the ASMR is associated with individual’s mood states rather than with his/her personality traits and that people with an anxious state tend to experience a strong ASMR. Our results provide important clues to understand the mechanisms of auditory–somatosensory interactions.

## Acknowledgements

This study was funded by JSPS KAKENHI grant (no. 17K04494 to H.M.K). We thank the Institute of Advanced Collaborative Research at Chukyo University for generous support.

